# Extracellular flux analysis and proteomic profiling indicate ibogaine promotes synaptic plasticity and neural repair by a metabolic preconditioning mechanism

**DOI:** 10.64898/2026.09.29.755437

**Authors:** Madison J. Oulton, Gracious D. S. Kasheke, Kelly A. Meaney, Robyn A. Aubie-Arseneault, Abhi Pushparaj, Jonathan E. Dickinson, Christopher S. Hughes, George S. Robertson

**Author notes:** Correspondence to: George S. Robertson, Department of Pharmacology, Faculty of Medicine, Dalhousie University, Halifax, Nova Scotia B3H 4R2, Canada.

## Abstract

We have examined how ibogaine may enhance synaptic plasticity and neural repair. Extracellular efflux analysis showed that ibogaine (3, 10 or 30 µM) produced a rapid (<5 minutes) and concentration-dependent suppression of mitochondrial function and glycolysis in mouse cortical neurons. Cell-based imaging demonstrated this metabolic inhibition was preceded by an elevation of the mitochondrial membrane potential and followed by increased mitochondrial reactive oxygen species levels. Like ibogaine, which blocks potassium channels, mitochondrial Kv1.3 channel (mitoKv1.3) inhibitors produce concordant changes over a similar timeframe. Following treatment with ibogaine (3, 10 or 30 µM) for 4 hours, a marked concentration-dependent rebound of energy production occurred 20 hours later suggestive of metabolic preconditioning. Indeed, proteomic profiling showed that oral ibogaine (100 mg/kg) administration to female mice produced changes in spinal cord protein abundance indicative of increasing metabolic stress from 0.25-24 hours that resolved by 72 hours. This was accompanied by activation, inhibition and reactivation of synaptic plasticity and neural repair signaling pathways at 0.25, 4-24 and 72 hours, respectively. Hence, by inducing transient metabolic stress, ibogaine may re-open a neurodevelopmental window characterized by enhanced synaptic plasticity and neural repair. Ibogaine binds with high affinity to the sigma-2 receptor. Like mitoKv1-3 inhibitors and sigma-2 receptor ligands, ibogaine transiently elevated AMP-activated protein kinase (0.25 hours) and temporally suppressed mechanistic target of rapamycin and lipid synthesis pathways (0.25-24 hours) implicated in enhanced synaptic plasticity and neural repair. These findings suggest ibogaine rewires and repairs neural circuits by a metabolic preconditioning mechanism, perhaps involving mitoKv1.3 and the sigma-2 receptor.

## 1. Introduction

Ibogaine is a hallucinogenic alkaloid derived from the root bark of a central African shrub called *Tabernanthe iboga* (Pope, 1969). This atypical psychedelic has been used for millennia by the Babongo pygmies, and more recently by practitioners of Bwiti and other traditional rites as part of “coming of age” ceremonies to induce an intense introspective dream-like state claimed to promote spiritual and mental well-being (de Rios et al., 2002; Fernandez, 1982). Since the 1980s, ibogaine has gained attention as a highly effective treatment for addiction (Davis et al., 2023; Mosca et al., 2023; Noller et al., 2018; Schenberg et al., 2014). More recently, ibogaine has been shown to dramatically reduce the symptoms of post-traumatic stress disorder in combat veterans suffering from head injuries (Cherian et al., 2024). Case reports also indicate that ibogaine alleviates neuropathic pain after brachial plexus nerve avulsion and motor deficits in multiple sclerosis (Chen et al., 2025a; Dickinson et al., 2023). These profound clinical improvements have been attributed to enhanced synaptic plasticity and neural repair (Biosca-Brull et al., 2024; Khan et al., 2021; Ly et al., 2018; Ona et al., 2023). However, the mechanisms are unclear.

Unlike neural cells, yeast do not have neurotransmitter receptors and transporters that are targeted by ibogaine (Mohammadi et al., 2015). However, like neural cells, yeast utilize mitochondria to produce energy (Lasserre et al., 2015). Studies in yeast have shown that ibogaine produces a burst of reactive oxygen species (ROS) production accompanied by a transient suppression of adenosine triphosphate (ATP) levels (Paskulin et al., 2010). Upon removal of ibogaine after treatment for 5 hours (hr), ATP concentrations rose rapidly in the yeast cultures to amounts well above basal levels (Paskulin et al., 2010). These findings suggest ibogaine produces mitochondrial dysfunction that triggers adaptations in response to the resultant metabolic stress. This hypothesis is supported by rodent studies that have shown ibogaine progressively elevates tricarboxylic acid (TCA) and glycolytic enzymes in the brain from 24 to 72 hr after administration (Paškulin et al., 2012). In terms of synaptic plasticity and neural repair, 24 hr after ibogaine administration brain-derived neurotrophic factor (BDNF) mRNA levels are reduced while at 72 hr they are dramatically increased in the prefrontal cortex of mice (Marton et al., 2019). BDNF production is closely coupled to mitochondrial function. Mitochondrial dysfunction lowers BDNF levels while enhanced mitochondrial function increases BDNF levels (Kim et al., 2015). These findings suggest ibogaine may regulate BDNF levels by altering mitochondrial function. In support of this metabolic hypothesis, dosing mice with the glycolysis inhibitor 2-deoxy glucose elevates BDNF expression by inducing an adaptive response to endoplasmic reticulum (ER) stress (Kumar et al., 2023).

Metabolic preconditioning of mitochondria is produced by exposing cells to mild, sub-lethal stressors (brief hypoxia, transient complex I or V inhibition using a low concentration of rotenone or Oligomycin A, respectively) that trigger adaptive pathways (Formentini et al., 2014; Ravati et al., 2001; Yuyun et al., 2013). This process is mediated by enhanced mitochondrial superoxide production which activates a wide array of signaling pathways (Shadel and Horvath, 2015). Tunicamycin, a potent inducer of ER stress, functions as a tool for metabolic preconditioning by inducing adaptive mechanisms between the ER and mitochondria. In its early stages, this stress increases physical coupling between the ER and mitochondria, promoting calcium transfer and enhancing mitochondrial bioenergetics (Bravo et al., 2011). Metabolic preconditioning enhances synaptic plasticity and neural repair by increasing BDNF levels, elevating mitochondrial function, and activating signaling pathways that regulate energy production, protein trafficking, lipid synthesis, transcription, translation, cytoskeletal dynamics and autophagy (Liang et al., 2021; Obrenovitch, 2008; Potter et al., 2010; Tu and Miyamoto, 2023). Based on these findings, we sought to determine if metabolic preconditioning pathways are engaged by ibogaine.

To gain insights into the effects of ibogaine on neuronal energy production, we first used XFe96 extracellular flux analysis to examine the concentration– and time-dependent effects of ibogaine on mitochondrial function and glycolysis in primary cultures of mouse cortical neurons. Using spinning disk confocal microscopy, we next measured the fluorescence signals generated by Tetramethylrhodamine, Methyl Ester, Perchlorate and mitoSOX Red in cortical neurons cultures treated with ibogaine to assess the concentration– and time-dependent effects of this atypical psychedelic on the mitochondrial membrane potential (Ψ_m_) and mitochondrial superoxide formation, respectively. Finally, we employed proteomic profiling to measure the abundance of 9,960 proteins in spinal cord samples of female mice 0.25, 4, 24 and 72 hr after an oral dose of ibogaine (100 mg/kg). Ingenuity pathway analysis (IPA) was used to reveal changes in the abundance of major functional classes of proteins. IPA suggested that ibogaine rewires and repairs the spinal cord by a metabolic preconditioning mechanism. Based on the results, of these studies, we propose molecular targets for ibogaine which may enable this atypical psychedelic to regulate the activities of vast signaling networks which re-open an early window in neurodevelopmental characterized by increased synaptic plasticity and neural repair.

## 2. Methods and materials

### 2.1. Animal Use

All animal experimentation in this project was approved by the University Committee on Laboratory Animals Ethics at Dalhousie University and procedures were conducted according to the Canadian Council on Animal Care guidelines. Female C57Bl/6 mice (20 g; Charles River) were housed in the Life Sciences Research Institute Animal Care Facility at Dalhousie University and had ad libitum access to food and water.

### 2.2. Primary cultures of mouse cortical neurons

Primary cortical neuron cultures were collected using E16 embryos from time-pregnant CD1 mice (Charles Rivers). Cells were plated at a density of 80,000 cells in 35 mm VWR confocal dishes, 1,000,000 cells/well in 6-well plates and 60,000 cells/well in 96-well Seahorse plates according to our previously described methods (Nichols et al., 2017). Media changes occurred on the fourth, seventh, tenth, twelfth, fourteenth, sixteenth and eighteenth day after plating, where half of the media was removed and replaced with fresh complete neurobasal media. Neuron cultures were used after 18-20 days in vitro (DIV) for all experiments. A stock concentration of ibogaine (0.1 mg/mL) were prepared by dissolving this compound in PBS. Ibogaine was added to the cultures in amounts sufficient to achieve concentrations of 3, 10 or 30 µM. Ibogaine was removed from the cell cultures at 4 hr by gently aspirating 75% of the Neurobasal media and replacing it with fresh media four times to lower the ibogaine concentration by 97%.

### 2.3. Primary cultures of mouse cortical astrocytes cultures

Astrocytes were isolated from cerebral cortices of CD1 pups (Charles River Canada, QC, Canada) at postnatal day 0 (P0) mice, as previously described (Novorolsky, 2023). Following isolation, astrocytes were maintained in astrocyte media [DMEM high glucose (pH = 7.4; Life Technologies, CA, USA) supplemented with 10% FBS (HyClone Laboratories Inc., UT, USA) and 20 µg/mL gentamycin (Thermo Fisher Scientific, MA, USA)] until they reached confluency (day in vitro 7-9). These cells were then passed and seeded at a density of 15,000 cells/well in 96-well Seahorse plates.

### 2.4. C8-B4 microglial cultures

The C8-B4 microglia cell line (ATCC, VA, USA) was maintained in Dulbecco’s Modified Eagle Medium (DMEM; pH = 7.4; Thermo Fisher Scientific, Cat. # 11995065, MA, USA). C8-B4 microglia were seeded at a density of 5,000 cells/well in 96-well Seahorse plates.

### 2.5. Seahorse XFe96 Extracellular Flux Analysis

Oxygen consumption rates (OCR) and extracellular acidification rates (ECAR) were measured using the Seahorse XFe96 extracellular flux analyzer (Agilent Technologies). The calibration plate used for sensor calibration was prepared and performed according to the manufacturer’s protocol. To allow the neurons to equilibrate to the assay conditions, the culture media was replaced with Seahorse XF assay medium [Dulbecco’s Modified Eagle’s Medium powder (8.3 g/L; Sigma-Aldrich) dissolved in Type 1 H_2_O, glucose (10 mM), sodium pyruvate (1 mM) and glutamine (2 mM)] and placed in a 37°C incubator 1 hr prior to running the assay. Mitochondrial function was examined by measuring OCR and ECAR at baseline and/or after the sequential addition of oligomycin (Oligo; 2 µM), carbonyl cyanide 4-(trifluoromethoxy) phenylhydrazone (FCCP; 1 µM), Rotenone (Rot; 300 nM), and Antimycin (5 µM). Three measurements were acquired at each condition for all experiments.

### 2.6. Reverse transcription quantitative polymerase chain reaction (RT-qPCR)

RNA was extracted from primary cortical neurons plated in 6-well plates using the Aurum total RNA minikit (Bio-Rad, cat# 732-6820) (according to the manufacturer’s instructions). Following RNA elution, the concentration and purity of RNA was estimated using the SPECTROstar Nano spectrophotometer (BMG Labtech). Absorbance readings (260 nm – 280 nm) above 1.8 were considered pure and included in the experiment. The iScript complementary DNA (cDNA) synthesis kit (Bio-Rad) was used to perform reverse transcription using 350 ng of template RNA from each sample. No template controls (no RNA template present) and no reverse transcriptase controls (without reverse transcriptase present) were run with every cDNA synthesis reaction. RT-qPCR was performed with the SsoFast EvaGreen Supermix kit (Bio-Rad). GAPDH, HPRT1 and OAZ1 were determined to be appropriate reference genes based on their collective stability using the BioRad CFX Maestro Software gene stability test. All RT-qPCR runs were performed with the Bio-Rad CFX96 real-time system C1000 touch thermal cycler. The RT-qPCR protocol conditions and melting curve program were completed as previously described with an annealing temp of 55°C (Nichols et al., 2017). Results were quality checked and analysed using the Bio-Rad CFX Maestro software. The resultant data were analysed using an ANOVA or non-parametric t-tests (Mann-Whitney) with PRISM (10.4.1) software. Target genes used in RT-qPCR analysis and the primer sequences are shown in Table 1 (Supplementary Information).

### 2.7. Fluorescence microscopy

To measure the Ψ_m_, neurons were incubated with Mitotracker Green (Thermofisher cat# M7514) at a concentration of 100 nM for 30 minutes (min) followed by gentle washing before incubation with Tetramethylrhodamine, Methyl Ester, Perchlorate (TMRM) (Thermofisher cat#T668) at a non-quenching concentration of 10 nM for 30 min. TMRM remained in solution for the duration of the experiment. Finally, neurons were incubated with Hoechst 33342 (Thermofisher cat#62249) at a concentration of 2 µM for 10 min. Oligo (4 µM) and FCCP (4 µM) were added at the end of the TMRM experiments. To evaluate mitochondrial superoxide production, neurons were incubated with MitoSOX Red ([6-(3,8-diamino-6-phenyl-5,6-dihydrophenanthridin-5-yl)hexyl]triphenylphosphonium iodide; Thermofisher cat# M36008) at a concentration of 1 µM for 30 min at 37°C. Next, the cells were washed gently with PBS and incubated with NucSpot^®^ Live 650 Nuclear Stain (Biotium #40082) for 10 min as per the manufacturer’s instructions. Images were captured on a Nikon CSU W1 SoRa Spinning disk confocal microscope using the 63X objective and analyzed using ImageJ analysis software.

### 2.8. Mass spectrometric detection of ibogaine

#### 2.8.1. Tissue preparation for mass spectrometric detection of ibogaine

Mice were euthanized 0.25, 4, or 24 hr following oral administration of ibogaine (100 mg/kg) by an overdose injection of sodium pentobarbital. Following loss of pedal withdrawal reflex, mice were intracardially perfused with PBS and spinal columns were dissected for further processing. Hydraulic extrusion of the spinal cord was then performed using a blunted 18-gauge needle attached to a 10 mL syringe filled with PBS. Spinal cord samples were supplemented with PBS at 1:4 (m:v) ratio, then samples were homogenized for 30 seconds (s) using 1.5 mm zirconium beads in a BeadBug^TM^ Microtube Homogenizer. Protein extraction was then performed cording to our previously described methods (Hughes et al., 2019).

#### 2.8.2 Mass Spectrometry Analysis

Targeted mass spectrometry analysis was carried out on a QTRAP5500 instrument interfaced to an Agilent 1290 Infinity II liquid chromatography (LC) system controlled using Analyst software (version 1.6.3). For analysis, samples were directly separated by an analytical column (2.1mm x 50mm, Kinetex F5 1.7µm, Phenomenex) at a flow rate of 0.4 mL/min. For analyte elution, the gradient held at an initial starting condition of 10% mobile phase B (99.9% acetonitrile, 0.1% formic acid) for 2-min before ramping to 60% B over 0.9 min, ramping to 90% B over 0.1-min and holding for 1-min, ramping to 10% B in 0.25-min, with a final equilibration at 10% B for 1.75-min (total run time of 6-min, mobile phase A = 0.1% formic acid in water). Data acquisition on the QTRAP5500 utilized a targeted selected reaction monitoring (SRM) acquisition scheme at ‘unit’ resolution with transitions for ibogaine (Q1 mass = 311.20, Q3 mass = 174.09, DP=120, CE=75, dwell=80msec), noribogaine (Q1 mass = 297.20, Q3 mass = 160.09, DP=120, CE=75, dwell=80msec), labeled lysine (Q1 mass = 155.10, Q3 mass = 137.10, DP=60, CE=12, dwell=20msec), and labeled threonine (Q1 = 125.10, Q3 = 60.10, DP=60, CE=21, dwell=20msec) with a total cycle time of 0.22 seconds. The QTRAP was globally set to use a positive ion spray voltage of 5500, CAD gas of 5, curtain gas of 25, temperature of 450, gas 1 of 55, and a gas 2 setting of 60. The entrance potential was globally set to 10, and the cell exit potential to 10. Retention times were around 3.14 min for ibogaine, 2.70 min for noribogaine, 0.29 min for lysine and 0.33 min for threonine.

### 2.9. Proteomic profiling

#### 2.9.1. Tissue preparation for proteomic profiling

Mice were euthanized 0.25, 4, 24, or 72 hr following oral administration of vehicle (150 µL) or ibogaine (100 mg/kg) by an overdose injection of sodium pentobarbital. Tissue preparation was completed as described above (section 2.5.1).

#### 2.9.2. Mass Spectrometry Analysis

Tissue lysates were prepared for proteomics by combining 50μL of disrupted tissue with 150 μL of a solution of 20 mM TrisCl pH 7.5, 150 mM KCl, 5 mM MgCl2, 0.5% (v/v) NP 40, 0.5% (w/v) sodium deoxycholate, and 0.5X Complete protease inhibitor (Sigma). To support degradation of DNA/RNA, 10 μL of a nuclease solution (1U/μL benzonase) was added to the solution and incubated at 24°C for 10-min. After nuclease digestion, 50μL of a solution of 600 mM TrisCl pH 7.5, 8% (w/v) SDS, and 10 mM dithiothreitol (DTT) was added to the tissue lysate and incubated at 95°C for 5-min in a heating block. After cooling to 24°C, lysates were alkylated using 40mM chloroacetamide (final concentration) and subsequently quenched with DTT. Protein concentration of the resulting lysate was measured using a BCA assay (ThermoFisher Scientific). Cleanup of extracted proteins prior to digestion was carried out using an adapted version of the previously described SP3 protocol (Hughes et al., 2014; Hughes et al., 2019). Specifically, 1 μL of a mixture of Sera-Mag carboxylate SpeedBeads (CAT#45152105050250 and CAT#65152105050250, prepared as 100 mg/mL stock solution in water) was added to 40 μg of protein for each sample prior to 4X volumes of acetone. Mixtures were incubated on a Thermomixer at 37°C for 5-min at 800rpm prior to centrifugation at 5,000 g for 5-min. After spinning, the supernatant was discarded and beads rinsed using 800 μL of an 80% (v/v) ethanol solution with pipetting to resuspend the beads. Mixtures were centrifuged at 5,000 g for 5-min and the supernatant discarded prior to the addition of 1 μg of ProteoSure trypsin mix (Marvelgent Biosciences) in 100 μL of digestion solution (100 mM ammonium bicarbonate, 1mM CaCl_2_) and incubation at 37°C for 16-hours in a Thermomixer with mixing at 800 rpm. After digestion, peptides were centrifuged at 12,000 g for 2-min and the supernatant recovered to a fresh tube containing 5 μL of a 10% (v/v) solution of trifluoroacetic acid. To desalt peptides prior to mass spectrometry (MS) analysis, an HPLC system (Agilent 1290 Infinity II with a DAD module) equipped with a reversed-phase column (CORTECS T3 1.6 μm, 2.1 x 50 mm, Waters) was utilized. Specifically, peptides were injected to a column equilibrated at 4% mobile phase B (0.1% formic acid in acetonitrile), rinsed for 1.5-min at a flow rate of 1mL/min, and subsequently eluted for 0.8-min at 80% mobile phase B (mobile phase A = 0.1% formic acid in water). Desalted peptides were dried in a SpeedVac centrifuge and reconstituted in 1% (v/v) formic acid in water at a concentration of 1 μg/μL based on the 210 nm UV signal from the HPLC desalting step.

Data independent acquisition (DIA) mass spectrometry (MS) analysis was carried out on an Orbitrap Lumos instrument (Thermo Scientific) interfaced to a Waters M-Class liquid chromatography (LC) system. For analysis, peptide samples were initially trapped on a pre-column (ACQUITY UPLC M-Class Symmetry C18 Trap, 180 µm x 2 cm, 5 µm beads, Waters) for 3-min at a flow rate of 20 µL/min of 1% mobile phase B (0.1% formic acid in acetonitrile) prior to separation by an analytical column (nanoEase M/Z Peptide BEH C18, 75 µm x 25 cm, 1.7 µm beads, Waters) coupled to a 20 μm inner diameter LOTUS electrospray tip (Fossil Ion Tech.). Gradient separation of trapped peptides started at an initial condition of 3% mobile phase B, ramped to 7% B in 1-min, to 28% B in 75-min, to 80% B in 0.5-min, hold at 80% B for 2-min, ramp to 3% B in 0.5-min, and a final hold at 3% B for 11-min for a total run time of 90-min (mobile phase A = 0.1% formic acid in water, flow rate = 300nL/min). For MS analysis on the Orbitrap Lumos, each peptide sample was injected three individual times (60-min total acquisition time for each), with each acquisition covering analysis of a separate mass range (injection 1 = 430 – 550, 2 = 550 – 690, 3 = 690 – 930 m/z). Specifically, the Orbitrap Lumos MS was globally set to use a positive ion spray voltage of 2000 V, an ion transfer tube temperature of 275°C, a default charge state of 3, and an RF Lens setting of 45%. The complete duty cycle of the acquisition method consisted of two MS1 scans and two sets of windowed MS2 scans (order MS1 – MS2 – MS1 – MS2). For the first, low mass range injection, the initial MS1 scan covered a mass range of 415-565 m/z at a resolution of 30,000 with an AGC target of 4e5 (100%) and a max injection time set to ‘Custom’ (54 milliseconds; ms). The following set of DIA MS2 scans covered a precursor range of 415-565 m/z with an isolation window size of 4 m/z (0 m/z overlap) for a total of 30 scan events. Each scan used an HCD energy of 30% and covered a defined mass range of 200-1800 m/z at a resolution of 30,000 with a normalized AGC target of 1000%, and the maximum injection time set to ‘Custom’ (54ms). Loop control was set to ‘N’ with a value of 30 spectra. The next MS1 scan and following DIA MS2 used the same settings as the previous with the exception that the MS2 precursor mass range was set to 428-552 m/z to give a 2 m/z stagger with the previous scan windows. For the second, medium mass range injection, the scan settings were the same as for the low mass injection with the following changes: 1. MS1 scan range of 535-705 m/z; 2. DIA-MS2 precursor scan ranges of 550-690 m/z and 548-692 m/z; 3. Loop control number of spectra = 35. For the third, high mass range injection, the scan settings were the same as the others with the following changes: 1. MS1 scan range of 675-945 m/z; 2. DIA-MS2 precursor scan ranges of 690-930m/z and 686-934 m/z; 3. DIA-MS2 isolation window of 8 m/z; 4. Loop control number of spectra = 30. All scan data were acquired in centroid mode. DIA-MS raw data were processed using DIA-NN (version 1.9.2) (Demichev et al., 2020). Specifically, a representative rat proteome fasta database (version 12/2024, 17,253 entries, includes contaminants) was provided to DIA-NN in order to perform an initial spectral library generation step [default settings for library generation, –-min-pr-mz 430 –– max-pr-mz 930 –-min-pr-charge 2 –-max-pr-charge 4, carbamidomethyl at cysteine as fixed, match between runs (MBR) disabled]. Afterwards, a subset of raw files for each mass range were searched together against this generated spectral library using DIA-NN with ‘unrelated runs’ activated to derive appropriate mass error and scan window settings. The entire set of raw riles for each mass range were then searched against the spectral library using the derived mass error and window settings with MBR enabled. Resulting report files were passed to the iq package (Lib.Q.Value = 0.01, Lib.PG.Q.Value = 0.01, Q.Value = 0.01, PG.Q.Value = 0.05) in R to generate estimates of protein abundance (Pham et al., 2020).

## 3. Results

### 3.1. Ibogaine produces a concentration-dependent suppression of mitochondrial function, glycolysis and BDNF mRNA levels in primary cortical neuron cultures

The selection of ibogaine concentrations for testing at 3, 10 and 30 µM in mouse cortical neuron cultures was based on literature values achieved in the brain after systemic injection of 40 mg/kg and our own mass spectrometric measurements of spinal cord concentrations after oral administration at dose of 100 mg/kg (Fig. S1). Addition of ibogaine (3, 10 and 30 µM) produced a concentration-dependent suppression of mitochondria function and glycolysis at 4 hr (Fig 1A and B). Basal respiration, ATP production, maximal respiration, spare respiratory capacity and proton leak were reduced in a concentration-dependent manner by ibogaine at 4 hr (Fig 1C-G). BDNF mRNA levels were also reduced in a concentration-dependent manner by ibogaine at 4 hr (Fig 1H).

**Fig 1.**
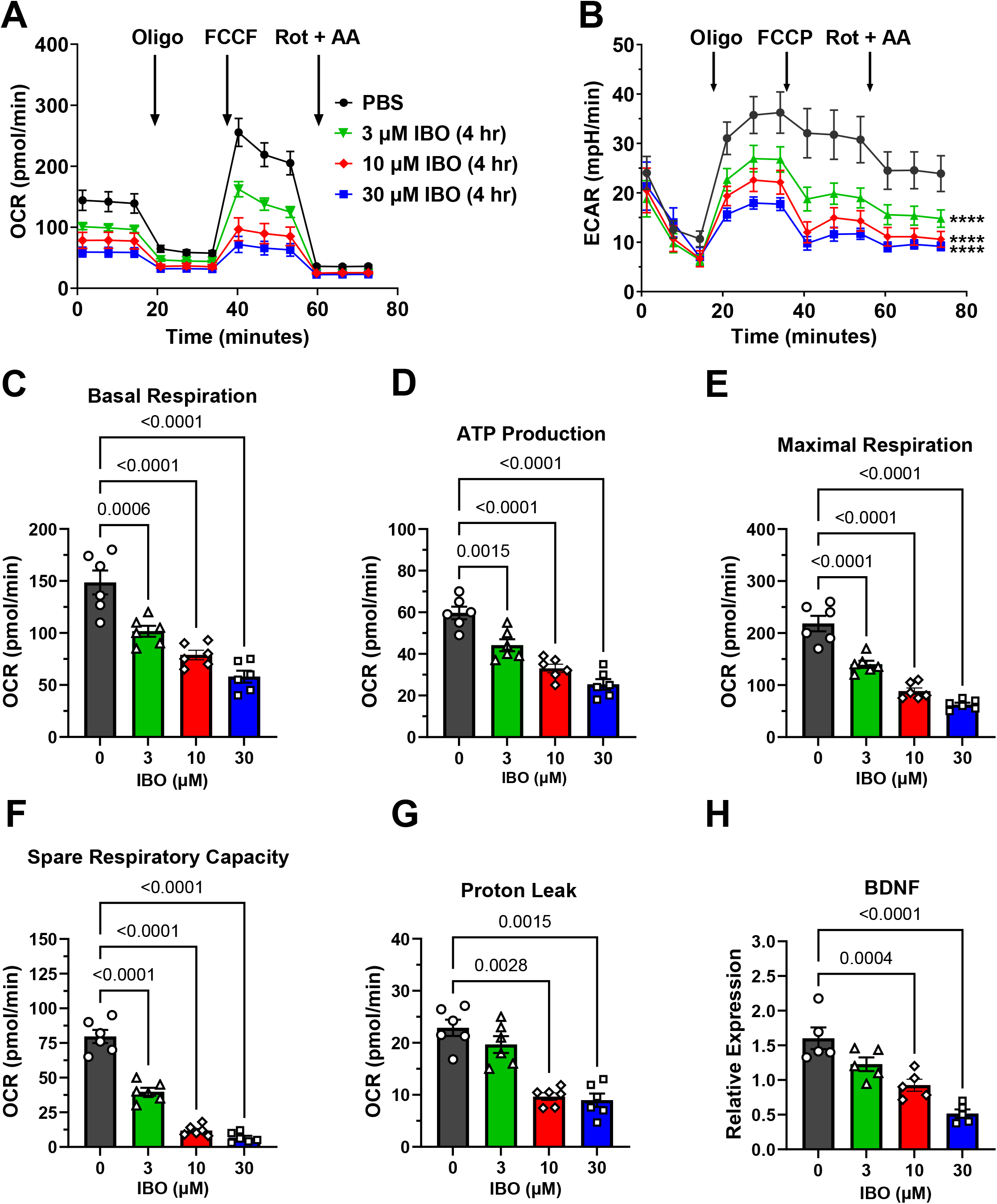
Oxygen consumption rates (OCR) (A), extracellular acidification rates (ECAR) (B), basal respiration (B), ATP production (C), maximal respiration (D), spare respiratory capacity (E), proton leak (F) and BDNF mRNA levels (G) in primary cortical neuron cultures 4 hr after treatment with ibogaine (3, 10 and 30 µM). Oligomycin (Olig), Carbonyl cyanide-p-trifluoromethoxyphenylhydrazone (FCCP), rotenone (Rot) and antimycin (AA). Data show the average valve ± SEM of quadruplicate measurements for 6 cell cultures.

### 3.2. Removal of ibogaine produces a concentration-dependent elevation of mitochondrial function, glycolysis and BDNF mRNA levels

Pharmacokinetic studies in rats (Hough et al., 1996; Rodrı Guez et al., 2020) as well as our own plasma (Fig. S1A) and spinal cord (Fig. S1B) measurements in female mice indicate that Ibogaine has an in vivo half-life of about 2 hr. To mimic the in vivo half-life of ibogaine, cortical neurons were treated with ibogaine (3, 10 or 30 µM) for 4 hr and then washed to remove ibogaine from the cultures. This treatment produced a marked elevation of mitochondrial function and glycolysis 20 hr later (24 hr; Fig 2A and B). Basal respiration, ATP production, maximal respiration, spare respiratory capacity and proton leak were increased in a concentration-dependent manner (Fig 2C-G). BDNF mRNA levels were also elevated in a concentration-dependent manner (Fig 2H).

**Fig 2.**
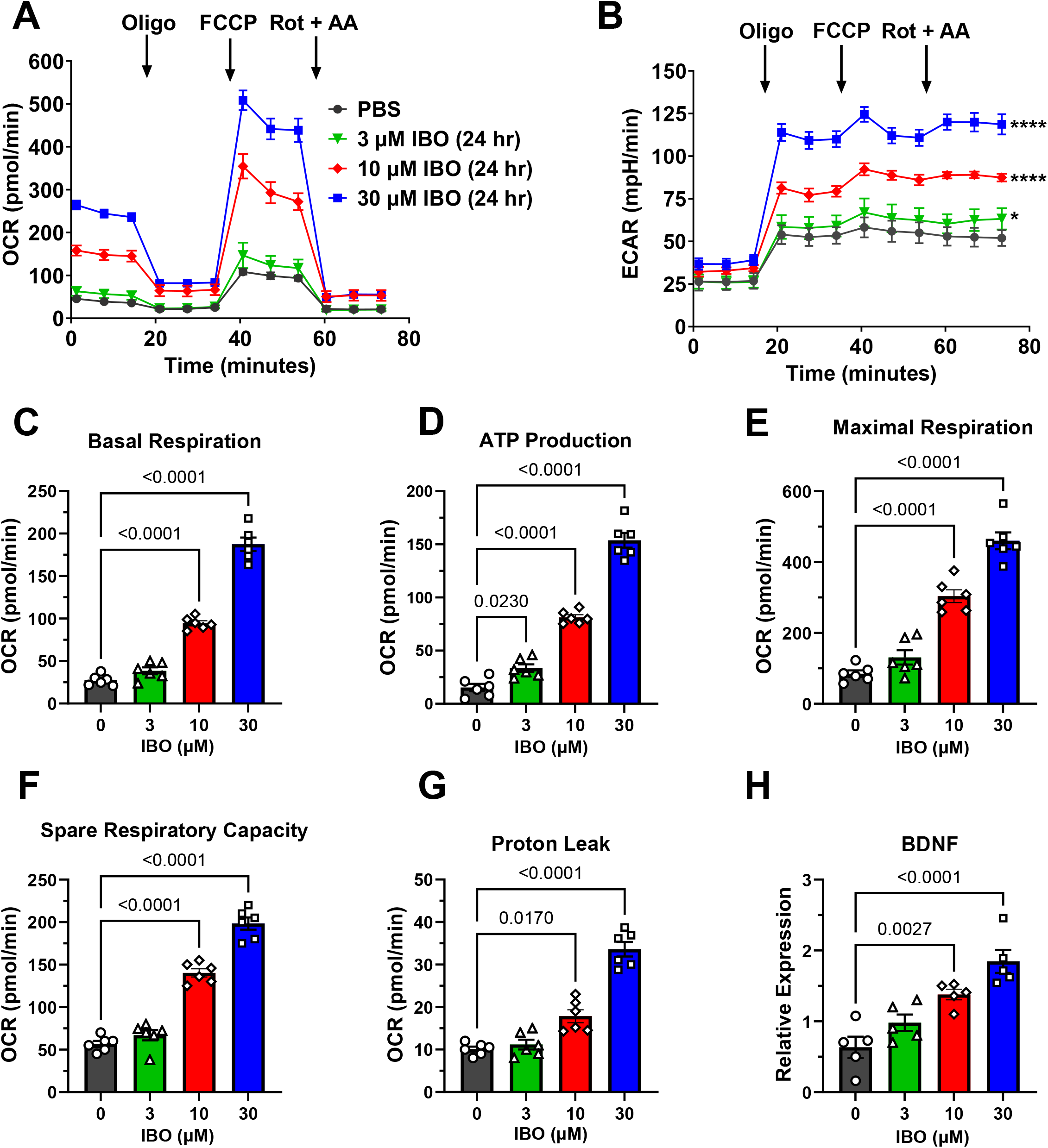
Oxygen consumption rates (OCR) (A), extracellular acidification rates (ECAR) (B), basal respiration (B), ATP production (C), maximal respiration (D), spare respiratory capacity (E), proton leak (F) and BDNF mRNA levels (G) in primary cortical neuron cultures 20 hr after treatment with ibogaine (3, 10 and 30 µM) for 4 hr (24 hr). Oligomycin (Olig), Carbonyl cyanide-p-trifluoromethoxyphenylhydrazone (FCCP), rotenone (Rot) and antimycin (AA). Data show the average valve ± SEM of quadruplicate measurements for 6 cell cultures.

### 3.3. Mitochondrial respiration and glycolysis are rapidly suppressed by the direct injection of ibogaine into the cortical neuron cell culture wells

To determine how rapidly ibogaine reduces cortical neuron mitochondrial respiration and glycolysis, we examined OCR and ECAR immediately after the delivery of ibogaine into the cell culture wells via the injection ports of the XFe96 instrument. Relative to control cultures, mitochondrial respiration (OCR) and glycolysis (ECAR) were rapidly suppressed (<5 min) in a concentration-dependent manner by the injection of ibogaine to generate well concentrations of 3, 10 or 30 µM (Fig 3A and B).

**Fig 3.**
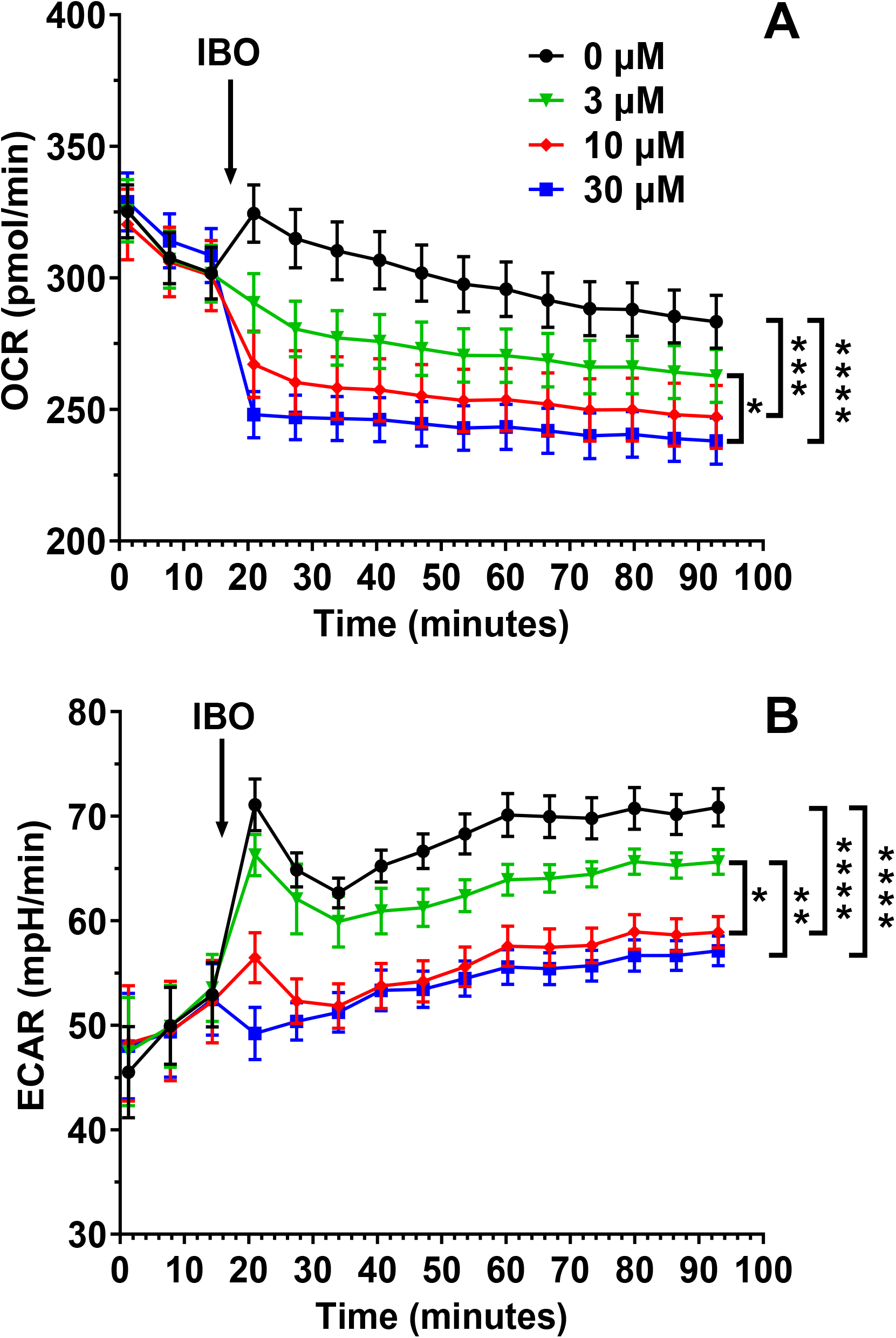
Oxygen consumption rates (OCR) (A), extracellular acidification rates (ECAR) (B) after port injections to achieve 3 or 10 µM concentrations of ibogaine. *p<0.05, **p<0.01, ***p<0.005. Data show the average valve ± SEM of quadruplicate measurements for 6 cell cultures.

### 3.4. Ibogaine produces a transient suppression of mitochondrial respiration and glycolysis in primary cortical astrocyte cultures and C8-B4 microglia

To investigate the effects of ibogaine on other neural cell subtypes, we measured mitochondrial respiration and glycolysis 1 and 4 hr after the addition of this compound to cultures of astrocytes (Fig. S2) or C8-B4 microglia (Fig 3S). Relative to control cultures, ibogaine (3 and 30 µM) produced a concentration-dependent suppression of OCR (Fig. S2A and 3SA) and ECAR (Fig. S2B and 3SB) at 1 hr in both astrocytes and microglia. By 4 hr, OCR (Fig. S2C and 3SC) and ECAR (Fig. S2D and 3SD) in astrocytes and microglia had returned to levels seen in control cultures.

### 3.5. Ibogaine produces a concentration-dependent elevation of the Ψ_m_ and mitochondrial ROS levels

We measured the fluorescence signals generated by TMRM and MitoSOX to evaluate whether ibogaine alters the Ψ_m_ and mitochondrial ROS levels, respectively. Within 2 min after the addition of 10 µM ibogaine, TMRM fluorescence intensity increased (Fig 4A and B) while mitochondrial ROS levels initially remained stable for 10 min (Fig 4D). Upon further incubation, ibogaine (3 and 10 µM) produced a concentration-dependent elevation of mitochondrial ROS levels at 40 and 60 min (Fig 4E).

**Fig 4.**
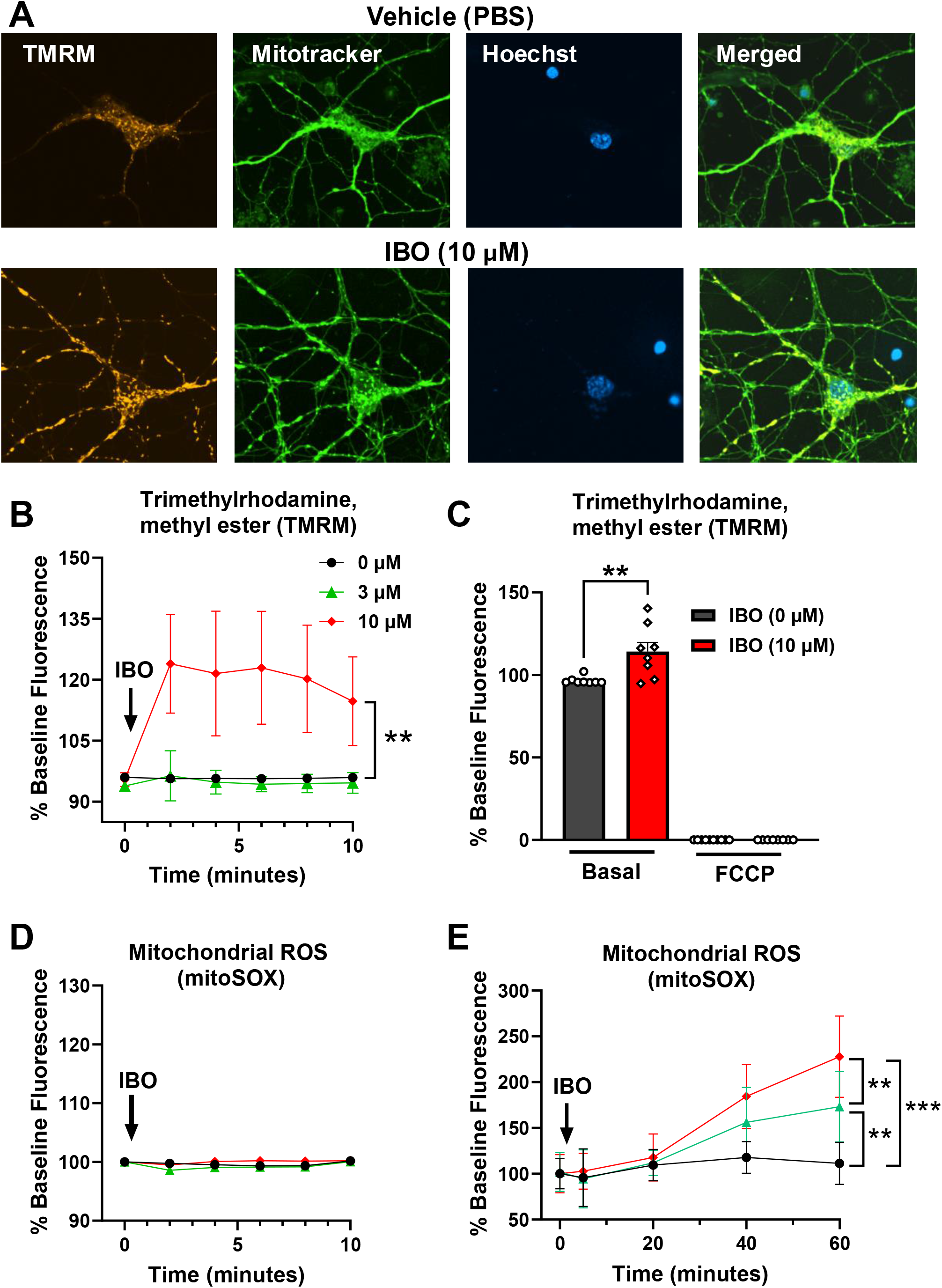
Primary cortical neurons stained with Tetramethylrhodamine, Methyl Ester, Perchlorate (TMRM), mitotracker green and Hoechst 33342. (A) and quantified over 10 min after treatment with ibogaine (3 and 10 µM) (B). Immediately following the experiment in B, FCCP was added directly to each well and fluorescence intensity quantified (C). Mitochondrial ROS (mitoSOX) was quantified after treatment with ibogaine (3 and 10 µM) for 10 min (D) or 60 min (E). *p<0.05, **p<0.01, ***p<0.005. Data show representative values for triplicate experiments.

### 3.6. Proteomic profiling revealed dynamic changes in the abundance of spinal cord proteins at varying time periods after the oral administration of ibogaine

Shotgun proteomics was used to measure the levels of 9,960 proteins in the spinal cord extracts of 6 female mice 0.25, 4, 24 and 72 hr after the oral administration of vehicle (150 µL) or ibogaine (100 mg/kg). Allometric scaling was used to select the oral dose of 100 mg/kg for mice (Reagan-Shaw et al., 2008) based on a flood dose of ibogaine (12 mg/kg) that alleviates the symptoms of post-traumatic stress disorder, neuropathic pain and multiple sclerosis (Chen et al., 2025a; Cherian et al., 2024; Dickinson et al., 2023). Relative to vehicle-treated mice, ibogaine reduced the levels of 1034, 1180, 1914 and 56 proteins and increased the levels of 1138, 800, 626, and 442 proteins at 0.25, 4, 24 and 72 hr, respectively (Fig. S4A-D).

### 3.7. Identification of changes in the abundance of major functional classes of proteins

IPA was performed to identify the metabolic and signaling networks regulated by ibogaine (Krämer et al., 2014). At 0.25 hr after the oral administration of ibogaine (100 mg/kg), protein abundance for 8 cellular stress pathways was increased (Fig 3A). These included the Coordinated Lysosomal Expression and Regulation (CLEAR) signaling, Kelch-like epichlorohydrin-associated protein 1 – Nuclear factor erythroid 2-related factor 2 (KEAP1-NFE2L2), production of reactive oxygen species (ROS), synthesis of ROS, nuclear factor erythroid 2-related factor 2 (NRF2)-mediated oxidative stress response, apoptosis, ER response and ultraviolet light UVA (315-400 nm)-induced mitogen-activated protein kinase signaling (UVA-induced MAPK signaling) pathways. Although the intra-Golgi and retrograde Golgi-to-ER traffic pathway was reduced, this decrease is also associated with increased cellular stress (Mohan et al., 2023). By 4 hr, a different pattern emerged. The CLEAR, apoptosis, ER response and UVA-induced MAPK signaling pathways remained elevated from 4-24 hr, but production of ROS was reduced at 4 hr and KEAP1-NFE2L2 and NRF2 oxidative stress response pathways were suppressed well below levels seen in vehicle-treated mice at 4 and 24 hr. By 72 hr, production of ROS and UVA-induced MAPK signaling returned to basal levels. By contrast, the CLEAR, apoptosis and ER stress response signaling pathways dropped below while the KEAP1-NFE2L2 and intra-Golgi and retrograde Golgi-to-ER traffic pathways rose to values above those for vehicle-treated mice.

In general, cellular metabolism pathways were suppressed from 0.25-24 hr by ibogaine (Fig 3B). These included concentration of lipids, synthesis of lipids, amino acids regulate mechanistic target of rapamycin (mTOR) signaling, oxidative phosphorylation, integration of energy metabolism, TCA cycle, cholesterol biosynthesis and glucose metabolism pathways. The only exception was the adenosine monophosphate-activated protein kinase (AMPK) signaling pathway which was elevated at 0.25 hr. By 72 hr, the abundance of proteins in the AMPK signaling, oxidative phosphorylation, TCA cycle, integration of energy metabolism, and cholesterol biosynthesis pathways had returned to basal levels. By contrast, concentration and synthesis of lipid, amino acids regulate mTORC1, mTOR signaling and glucose metabolism pathways increased to levels above those for vehicle-treated mice.

IPA detected dynamic changes for 10 RNA and translation pathways (Fig 4A). At 0.25 hr, mitochondrial translation, splicing of RNA, microRNA biogenesis signaling pathway, RNA polymerase II transcription and processing of capped intron containing pre-mRNA pathways were elevated while major pathway of rRNA processing in the nucleolus and cytosol, nonsense-mediated decay, eukaryotic translation initiation, Eukaryotic Initiation Factor 2 (EIF2) signaling and the ribosomal quality control pathway were depressed by ibogaine. These trends remained evident at 4-24 hr except for the microRNA biogenesis signaling and processing of capped intron containing pre-mRNA pathways that were depressed. The only difference from these trends was that mitochondrial translation pathway was depressed at 24 hr. By 72 hr, all pathways had returned to basal levels except for the RNA polymerase II transcription and processing of capped intron containing pre-mRNA pathways that were elevated with respect to vehicle-treated mice.

IPA revealed changes in 9 protein degradation and clearance pathways (Fig 4B). At 0.25 hr, macroautophagy was elevated while fusion and formation of lysosomes, deubiquitination, autophagy, protein ubiquitination pathway, microautophagy signaling pathway and neddylation pathways were suppressed. Except for the sumoylation pathway that was elevated, these trends remained unchanged from 4-24 hr. At 72 hr, fusion and formation of lysosomes and the sumoylation pathways returned to basal levels while macroautophagy, deubiquitination, autophagy protein ubiquitination, microautophagy signaling and neddylation pathways reversed directions becoming higher than vehicle-treated mice.

IPA identified changes in 11 synaptic plasticity pathways (Fig 5A). At 0.25 hr, ephrin receptor signaling, microtubule dynamics, reelin signaling in neurons, signaling by neurogenic locus notch homolog protein 1 (NOTCH1) and the synaptogenesis signaling pathways were elevated by ibogaine. By contrast, these pathways were depressed at 4 and 24 hr. From 0.25-24 hr, signaling by Roundabout (ROBO) receptors, degradation of beta-catenin by the destruction complex, Ras homologous (RHO) guanosine triphosphatase (GTPase) cycle, cyclin-dependent kinase 5 (CDK5) and T-cell Factor (TCF) dependent signaling in response to wingless-related integration site (WNT) pathways were increasingly suppressed. By 72 hr, protein abundance for all 11 synaptic plasticity signaling pathways was increased relative to vehicle-treated mice.

**Fig 5.**
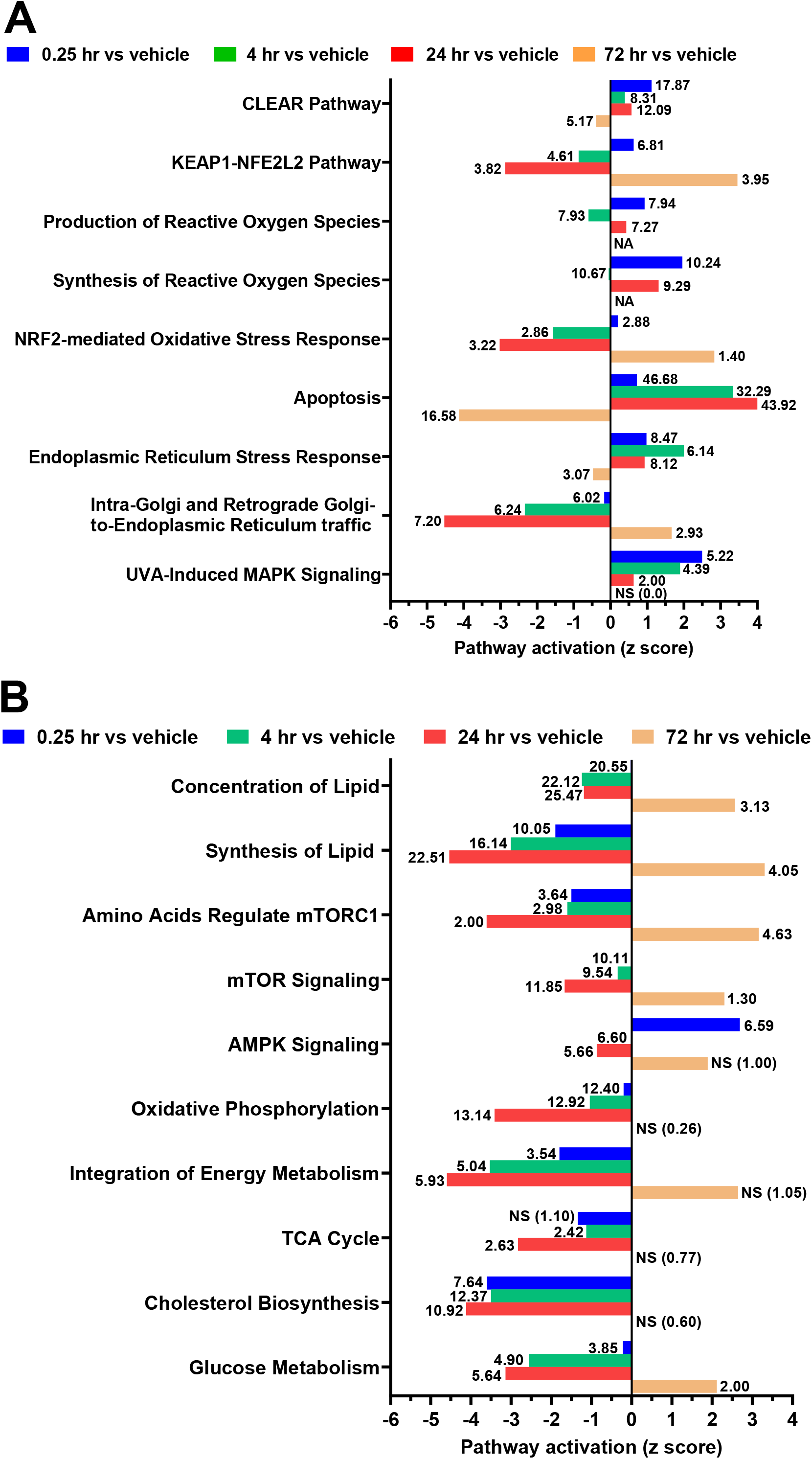
Z scores for cellular stress. (A) and energy (B) signaling pathways in the spinal cord based on data for 6 female mice 0.25, 4, 24 and 72 hr after an oral dose of ibogaine (100 mg/kg). The –logP values are indicated for each pathway. Values less than 1.3 were considered non-significant (NS). Not available (NA).

**Fig 6.**
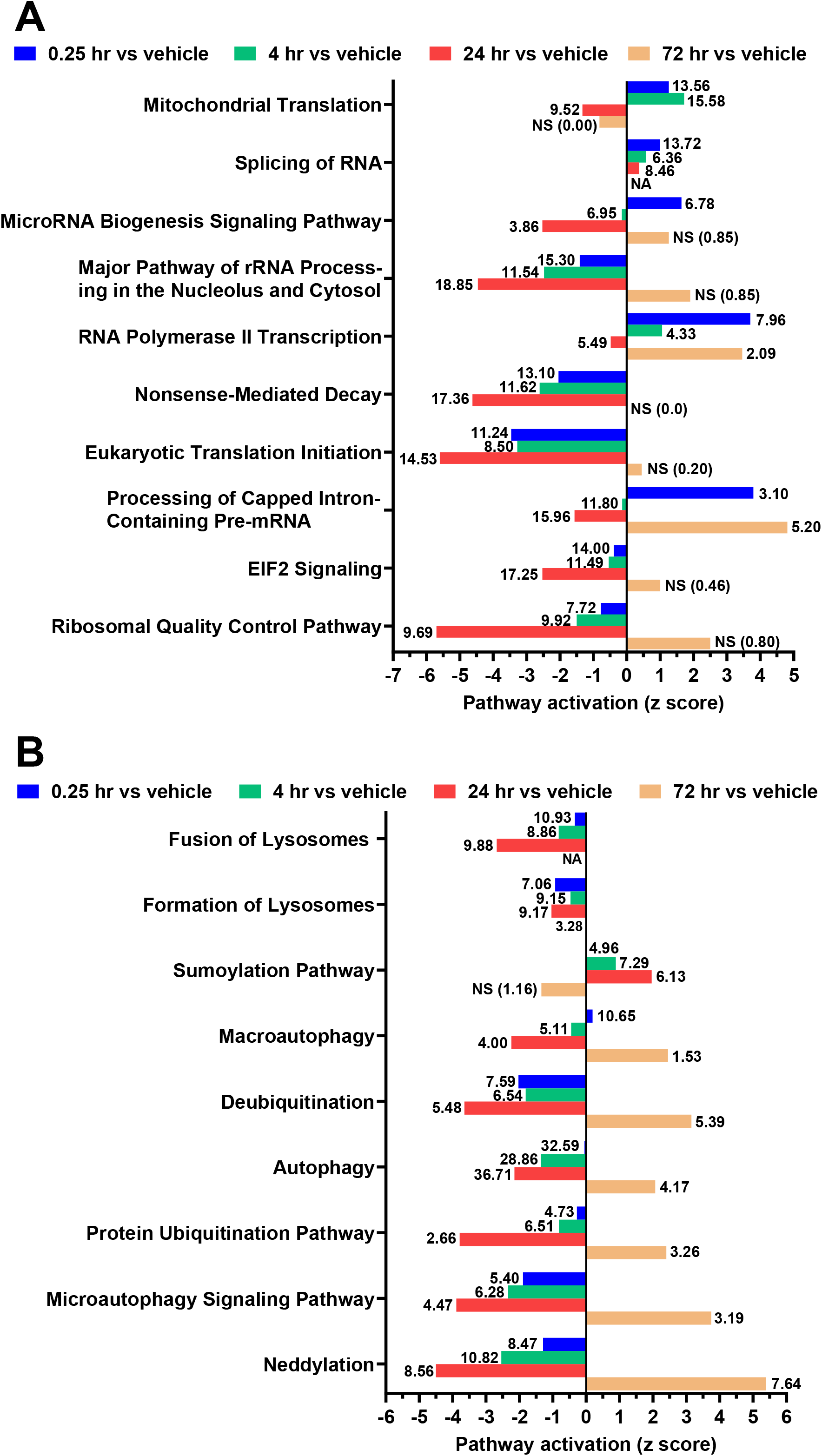
Z scores for RNA processing and translation. (A) and lipid level and protein degradation (B) signaling pathways in the spinal cord based on data for 6 female mice 0.25, 4, 24 and 72 hr after an oral dose of ibogaine (100 mg/kg). The –logP values are indicated for each pathway. Values less than 1.3 were considered non-significant (NS). Not available (NA).

**Fig 7.**
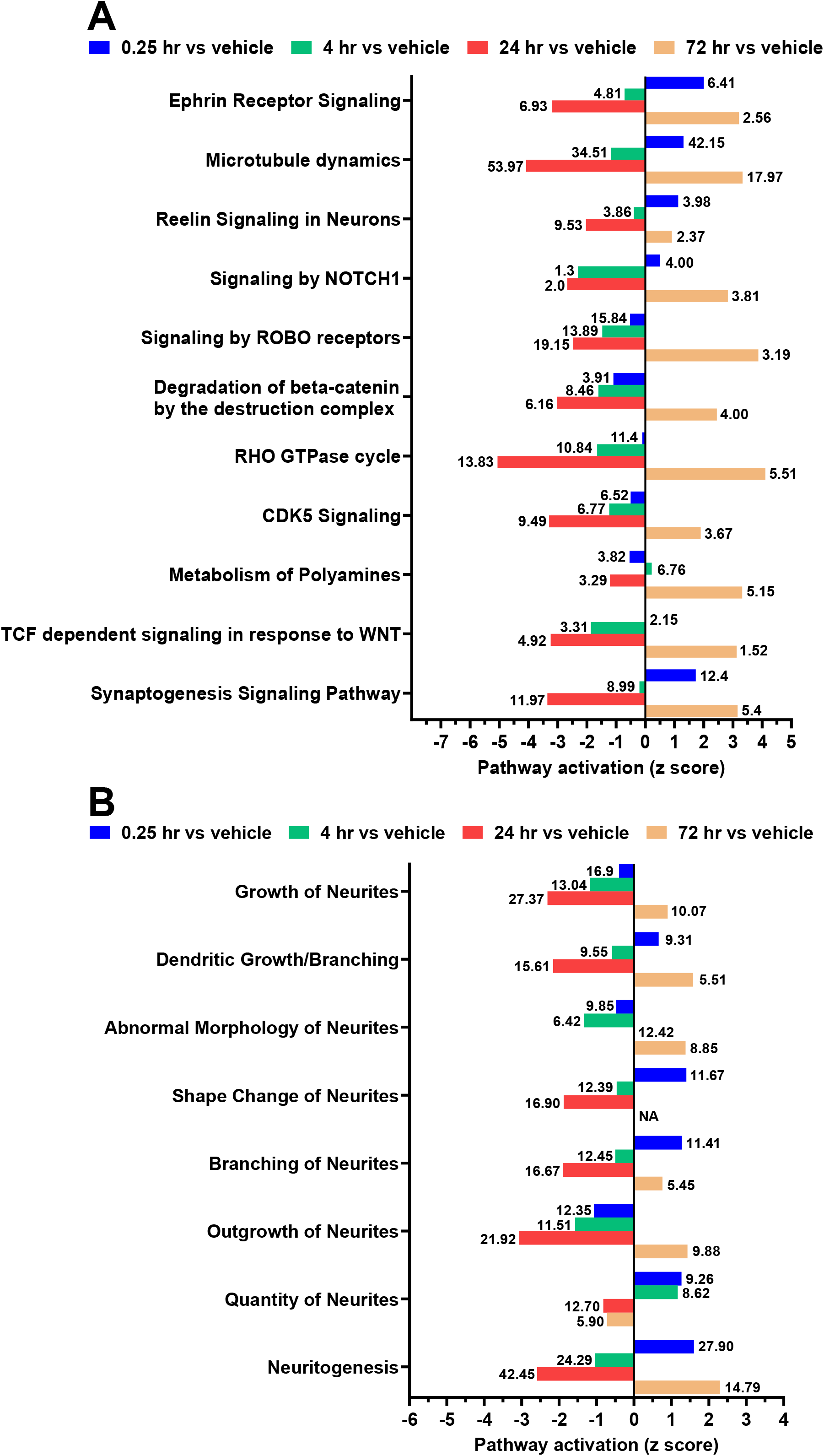
Z scores for synaptic plasticity. (A) and neurite (B) signaling pathways in the spinal cord based on data for 6 female mice 0.25, 4, 24 and 72 hr after an oral dose of ibogaine (100 mg/kg). The –logP values are indicated for each pathway. Values less than 1.3 were considered non-significant (NS). Not available (NA).

IPA detected changes in 8 neurite pathways (Fig 5B). At 0.25 hr, growth of neurites and abnormal morphology of neurites were reduced while dendritic growth/branching, shape change of neurites, branching of neurites, quantity of neurites and neuritogenesis pathways were increased by ibogaine. From 4-24 hr, dendritic growth/branching, shape change of neurites, branching of neurites and neuritogenesis pathways declined to levels below those for vehicle-treated mice. By contrast, there was a highly significant upregulation of growth of neurites, dendritic growth/branching, abnormal morphology of neurites, branching of neurites and neuritogenesis pathways at 72 hr. The shape change of neurites pathways returned to basal levels at 72 hr.

## 4. Discussion

We first used XFe96 extracellular flux analysis to examine the concentration– and time-dependent effects of ibogaine on mitochondrial function and glycolysis in primary cultures of mouse cortical neurons and astrocytes and the C8-B4 microglial cell. In all cases, ibogaine produced a concentration-dependent suppression of mitochondrial function and glycolysis. By comparison to neurons, these decreases were of smaller magnitude at 1 hr and disappeared at 4 hr in astrocytes and microglia. Port injections indicated ibogaine caused a rapid (<5 min) and concentration-dependent reduction in neuronal energy production. Removal of ibogaine from the cortical neuron cultures at 4 hr resulted in marked concentration-dependent increases in mitochondrial function and glycolysis at 24 hr. These findings are in keeping with yeast studies in which ibogaine (3, 12 and 60 µM) produced a concentration-dependent reduction of ATP levels (Paskulin et al., 2010). This fall in ATP levels was attributed to increased ATP consumption that depleted the pool of ATP (Paskulin et al., 2010). To assess the effects of ibogaine on ATP synthesis, we measured OCR after the inhibition of ATP synthase with oligomycin. These measurements suggest ibogaine may also deplete ATP levels in neurons and astrocytes by reducing ATP synthesis. However, based just on the effects of oligomycin on OCR, we cannot rule out the possibility that ibogaine deletes the pool of ATP by increasing ATP consumption.

Hyperpolarization of the Ψ_m_ produces an immediate elevation of mitochondrial ROS production (Zorov et al., 2006). Hence, the large delay between Ψ_m_ hyperpolarization and increased mitoSOX fluorescence indicates a non-mitochondrial trigger for increased mitochondrial ROS levels. Our proteomic profiling studies show oral ibogaine administration rapidly (0.25 hr) activates the ER stress response pathway in the spinal cord. The Ψ_m_ hyperpolarizes quickly as part of an adaptive response to ER stress that is followed by a gradual increase in mitochondrial ROS levels (Chen et al., 2025b; Knupp et al., 2019). Hence, ER stress may be responsible for the delayed rise in mitoSOX fluorescence after ibogaine treatment.

Ibogaine and its primary metabolite noribogaine are thought to produce cardiac arrhythmias by blocking human ERG (hERG) potassium channels in the heart (Koenig et al., 2014; Thurner et al., 2014). The highly lipophilic nature of ibogaine may also indirectly inhibit the hERG channel as well as other potassium channels such as mitoKv1.3 by altering the local lipid microenvironment (Bock et al., 2003; Teisseyre et al., 2021; Zünkler, 2024). Compounds that block mitoKv1.3 hyperpolarize the Ψ_m_ by suppressing potassium ion movement from the cytosol to the mitochondrial matrix (Szabo et al., 2008). Without positive potassium ions entering the matrix to counter-balance proton pumping, the electrical gradient between the intramembrane space and matrix becomes excessively negative (Szewczyk, 2024). As a result, respiratory complexes (I, III, and IV) cannot pump protons against this elevated electrical barrier, forcing electron transport to halt and causing a sharp drop in mitochondrial respiration (Wikström and Springett, 2020). Electrons therefore escape from the respiratory chain which directly reduce molecular oxygen resulting in elevated ROS formation (Larosa and Remacle, 2018). Blockade of mitoKv1.3 activity with small molecule inhibitors hyperpolarizes the Ψ_m_(Szabo et al., 2008), lowers mitochondrial respiration and increases mitochondrial ROS production (Leanza et al., 2017; Styles et al., 2021) over a similar period as ibogaine. Finally, like ibogaine, MitoKv1.3 inhibition has been shown to suppress glycolysis (Prosdocimi et al., 2024).

Proton leak is defined as the movement of protons from the intramembrane space to the matrix which is not coupled to ATP synthesis (Chance and Williams, 1955). Increased proton leak attenuates electron escape from the respiratory chain responsible for the generation of ROS (Divakaruni and Brand, 2011). By contrast, reduced proton leak promotes mitochondrial ROS production (Divakaruni and Brand, 2011). Hence, our findings suggest that incubation with ibogaine for 4 hr increased mitochondrial ROS levels by reducing proton leak. Conversely, increased proton leak 20 hr after the removal of ibogaine (24 hr) may have reduced mitochondrial ROS levels. Mitochondrial function and BDNF expression are closely linked (Cheng et al., 2010). Treatments that suppress mitochondrial function reduce BDNF mRNA levels while increased mitochondrial function promotes BDNF expression (Liu et al., 2025). It is therefore tempting to speculate that suppressed followed by enhanced mitochondrial function mediates reduced and elevated BDNF mRNA levels by ibogaine at 4 and 24 hr, respectively.

Mass spectrometry was employed to measure the levels of 9,960 proteins in spinal cord extracts of female mice 0.25, 4, 24 and 72 hr after the oral administration of vehicle or ibogaine (100 mg/kg). Allometric scaling was used to select the oral dose of 100 mg/kg for mice (Reagan-Shaw et al., 2008) based on a flood dose of ibogaine (12 mg/kg) that alleviates the symptoms of post-traumatic stress disorder, neuropathic pain and multiple sclerosis (Chen et al., 2025a; Cherian et al., 2024; Dickinson et al., 2023). IPA was then performed to identify the metabolic and signaling networks regulated by ibogaine. In keeping with impaired mitochondrial function in cortical neurons treated with ibogaine for 4 hr, ibogaine induced a cellular stress response characterized by elevated CLEAR, KEAP-NFE2L2, synthesis and production of ROS, NRF2-mediated oxidative stress response, apoptosis, ER stress and UVA-induced MAPK signaling and suppression of intra-Golgi and retrograde Golgi-to-ER trafficking pathways at 0.25 hr. Cellular stress causes an accumulation of unfolded proteins in the ER that activates the CLEAR and apoptosis signaling and depresses intra-Golgi and retrograde Golgi-to-ER trafficking pathways (Ballabio and Bonifacino, 2020; Mohan et al., 2023; Ray et al., 2012; Shirai and Yamauchi, 2024). Beginning at 4 hr, adaptations to cellular stress became evident as suggested by the decline of KEAP-NFE2L2, production and synthesis of ROS and NRF2-mediated oxidative stress response signaling pathways that peaked at 24 hr. Consistent with a rebound elevation of mitochondrial function after removal of ibogaine from the cortical neuron cultures, KEAP-NFE2L2, NRF2-mediated oxidative stress response and intra-Golgi and retrograde Golgi-to-ER trafficking pathways increased above basal levels while the CLEAR and apoptosis signaling pathways fell below them at 72 hr. Consistent with evidence that these changes oppose cellular stress (Sasaki and Yoshida, 2015), the ER stress response and UVA-induced MAPK signaling pathways were reduced at 72 hr.

IPA revealed that ibogaine had profound effects on cellular metabolism in the spinal cord. Among the most prominent changes were markedly increased AMPK signaling at 0.25 hr, reduced lipid and cholesterol synthesis, amino acids that regulate mTOR, mTOR signaling, oxidative phosphorylation, integration of energy metabolism, TCA cycle and glucose metabolism pathways from 0.25-24 hr. These changes reflect a metabolic block where the cell prioritizes immediate energy conservation over building complex molecules to promote survival (Krycer et al., 2020; Long and Zierath, 2006; Ricoult and Manning, 2013). Elevation of the ER stress response and suppression of intra-Golgi and retrograde Golgi-to-ER traffic pathways also disrupt cholesterol synthesis by blocking transport of sterol regulatory element-binding protein transcriptional factors from the ER to the Golgi apparatus (Xu et al., 2015). A major cause of impaired Golgi trafficking is lysosomal dysfunction that causes an accumulation of undegraded material and failure of organelle-to-organelle signaling (Settembre and Perera, 2024). In support of this lysosomal mechanism, both fusion and formation of lysosome pathways were reduced by ibogaine from 0.25-24 hr.

The induction of transient cellular stress by ibogaine was also indicated by changes in the abundance of proteins for RNA processing and translation pathways. Inhibition of eukaryotic translation initiation, EIF2 signaling and the ribosomal quality control pathways reflect induction of the unfolded protein response that suppresses protein translation to reduce ER load (Guan et al., 2014). Increased processing of capped intron-containing pre-mRNA is an adaptive response to ER stress that rapidly elevates the production of mature mRNA from a pre-existing pool of transcripts (Rambout and Maquat, 2024). Reduced nonsense-mediated mRNA decay facilitates this process by stabilizing transcripts necessary for a robust response to cellular stress (Hug et al., 2016). Increased splicing of RNA is a hallmark of ER stress that selectively accelerates the processing of key mRNAs to expand cellular protein-folding capacity (Yoshida et al., 2001). Suppression of the major pathway of rRNA processing in the nucleolus and cytosol acts as a “pause” mechanism to conserve energy and maintain cellular integrity during metabolic stress (Szaflarski et al., 2022). The elevation of RNA polymerase II transcription supports induction of microRNA biogenesis that is an energy efficient mechanism for fine tuning the adaptive response to cellular stress (Byrd and Brewer, 2013).

IPA detected a general suppression of ubiquitination, neddylation and autophagy networks from 0.25-24 hr. Our findings indicate this occurs because mitochondrial dysfunction blocks these energy-dependent processes by depleting ATP and generating high levels of ROS which inhibit ubiquitin and neddylation activating ligases and proteasomal function (Karbowski et al., 2022). The subsequent buildup of damaged proteins may thus exacerbate ER stress. Since enhanced AMPK signaling stimulates sumoylation (Seager et al., 2024) the marked induction of AMPK signaling at 0.25 hr may reflect increased sumoylation. Finally, the profound rebound increases in microautophagy, macroautophagy, neddylation, deubiquitination, autophagy and protein ubiquitination pathways at 72 hr may promote synaptic plasticity and neural repair by clearing damaged cellular components, degrading axonal repulsive proteins and stabilizing pro-survival factors (Grabbe et al., 2011; Haas and Broadie, 2008; Vucic et al., 2011).

IPA detected elevated ephrin receptor signaling, microtubule dynamics, reelin signaling in neurons, signaling by NOTCH1 and the synaptogenesis signaling pathways at 0.25 hr. The protein abundance for these pathways was progressively reduced from 4-24 hr to below basal levels. The transient increase of these pathways suggests ibogaine produced a burst of synaptic remodeling at 0.25 hr (Alberi et al., 2011; Cingolani and Goda, 2008; Hethorn et al., 2015; Jaworski et al., 2009; Rasool and Jahani-Asl, 2024). Signaling by ROBO receptors, degradation of beta-catenin by the destruction complex, RHO GTPase cycle, CDK5, TCF dependent signaling in response to WNT and metabolism of polyamines were increasingly suppressed from 0.25-24 hr. At 72 hr there was a rebound elevation of all synaptic plasticity signaling pathways suggestive of the formation and strengthening of new connections (Blockus et al., 2021; Sagar et al., 2021; Tang, 2014; Tolias et al., 2011). Of relevance to the mechanism of action of ibogaine in the treatment of pain after nerve injury, reduced ephrin receptor, NOTCH1 and WNT signaling in the spinal cord alleviates mechanical allodynia and thermal hyperalgesia in rodent models of neuropathic pain (Battaglia et al., 2003; Tang et al., 2022; Xie et al., 2015).

IPA provided key insights into how ibogaine influences neurite pathways. At 0.25 hr, shape change, branching and quantity of neurites pathways as well as dendritic growth/branching and neuritogenesis signaling pathways were elevated. Like many of the previously described metabolic and signaling networks, these neurite pathways decreased from 4-24 hr and then rebounded above basal levels and 72 hr. This up-down-up cycle matches the outgrowth, stability/pruning and regrowth/adaptive remodeling stages of synaptic plasticity during neurodevelopment (Bianchi et al., 2013; Faust et al., 2021; Gazerani, 2025; Hruska et al., 2015). Indeed, ibogaine administration has been reported to induce juvenile-like synaptic plasticity in the visual cortex of adult mice (Acuña et al., 2025). Interestingly, the highly potent and preferential mKv1.3 inhibitor Psora-4 has been shown to increase the differentiation of neural progenitor cells into electrically active neurons (Zhou et al., 2015). These findings suggest that ibogaine re-opens a developmental window characterized by enhanced synaptic plasticity and neural repair that may be mediated in part by mKv1.3 inhibition.

The profound reductions in glucose metabolism, antioxidant, cholesterol and lipid synthesis, mTOR, RNA processing, ubiquitination, neddylation, microautophagy, autophagy, ROBO receptor, beta-catenin and Wnt signaling pathways from 0.25-24 hr followed by a rebound elevation at 72 hr are adaptations to ER stress (Honda et al., 2005; Röhrl et al., 2014; Sheng et al., 2023; Yang et al., 2024). The induction of these processes supports synaptic plasticity and neural repair by enhancing antioxidant defense, clearing damaged lipids and proteins, remodeling dendrites and neurites and fueling the production of energy and metabolic substrates for myelination, axonal growth and synaptogenesis (Berghoff et al., 2022; Cantuti-Castelvetri et al., 2018; Zhang et al., 2020). In these regards, it is worth noting that the active metabolite noribogaine potently activates the kappa opioid receptor (EC_50_=0.61 µM) known to increase myelin repair (Maillet et al., 2015). Hence, future studies should examine whether ibogaine and noribogaine have complimentary effects that enhance CNS repair. The high affinity of ibogaine for the ER and lysosome-associated chaperone protein TMEM97 may contribute to the rapid and profound suppression of cholesterol synthesis produced by ibogaine (Alon et al., 2017; Bowen et al., 1995). This chaperone protein forms a complex with progesterone membrane binding component-1 and the low-density lipid receptor that regulates cholesterol trafficking and synthesis (Ebrahimi-Fakhari et al., 2016; Riad et al., 2018). TMEM97 ligands have been shown to induce mild ER stress by overloading the lysosome with cholesterol that triggers a rapid inhibition of cholesterol synthesis (Takchi et al., 2024). Future studies that employ conditional and cell-specific TMEM97 knockout mice (Hong et al., 2024) as well as mass spectrometric techniques to measure the metabolome and identify ibogaine binding partners (Li et al., 2025) should prove useful in understanding the molecular basis for the therapeutic actions of ibogaine.

## FIGURE LEGENDS

**Fig S1.** Plasma and spinal cord concentrations of ibogaine 0.25, 1, 4 and 24 hr after an oral dose of ibogaine (100 mg/kg). Data show the average ± SEM for 4 female mice.

**Fig S2.** Oxygen consumption rates (OCR) and extracellular acidification rates (ECAR) in primary cortical astrocyte cultures 1 (A and B) and 4 (C and D) hr after treatment with ibogaine (3 or 30 µM). Oligomycin (Olig), Carbonyl cyanide-p-trifluoromethoxyphenylhydrazone (FCCP), rotenone (Rot) and antimycin (AA). Data show the average valve ± SEM of triplicate measurements for 6 cell cultures. *p<0.05, **p<0.01, ***p<0.005 relative to PBS control for all other groups.

**Fig S3.** Oxygen consumption rates (OCR) and extracellular acidification rates (ECAR) in C8-B4 microglia 1 (A and B) and 4 (C and D) hr after treatment with ibogaine (3 and 30 µM). Oligomycin (Olig), Carbonyl cyanide-p-trifluoromethoxyphenylhydrazone (FCCP), rotenone (Rot) and antimycin (AA). Data show the average valve ± SEM of triplicate measurements for 6 cell cultures. *p<0.05, **p<0.01, ***p<0.005 relative to PBS control for all other groups.

**Fig S4.** Volcano plots showing the distribution of down and upregulated proteins in the spinal cord 0.25 hr (A). 4 hr (B), 24 hr (C) and 72 hr (D) after an oral dose of ibogaine (100 mg/kg). Each point represents the average value for 6 female mice.

## DATA AVAILABILITY

The mass spectrometry proteomics data have been deposited to the ProteomeXchange Consortium via the PRIDE [1] partner repository with the dataset identifier PXD082316. The data for the mass spectrometrometric detection of ibogaine are available in the <u>Mendeley Data</u> repository, at https://data.mendeley.com.

All other data are available upon request.

## AUTHOR CONTRIBUTIONS

MJO and KAM performed the XFe96 extracellular efflux analysis and related data analysis. RAA-A performed the RT-qPCR and related data analysis. MJO performed fluorescence microscopy and related data analysis. CSH performed the mass spectrometry. GDSK performed the proteomic profiling and related data analysis using IPA. AP and JED assisted with the manuscript preparation. GSR assisted with all data analysis and manuscript preparation.

## FUNDING

This work was funded by research grants from the MITACS Accelerate program (GSR; project number IT39741) and the Department of Psychaitry Research Fund at Dalhousie University.

## ACKNOWLEDGEMENTS

We thank Drs. Anatoly Starkov and Evgeny Pavlov (Department of Molecular Pathobiology, New York University) for useful discussions about the effects of ibogaine on mitochondrial function.

## COMPETING INTERESTS

AP and JED are affiliated with Ambio Life Sciences which offers ibogaine treatments. JED is a shareholder in Ambio Life Sciences. JED is a shareholder of Terragnosis Inc. which manufactures and distributes ibogaine. The remaining authors have nothing to disclose.

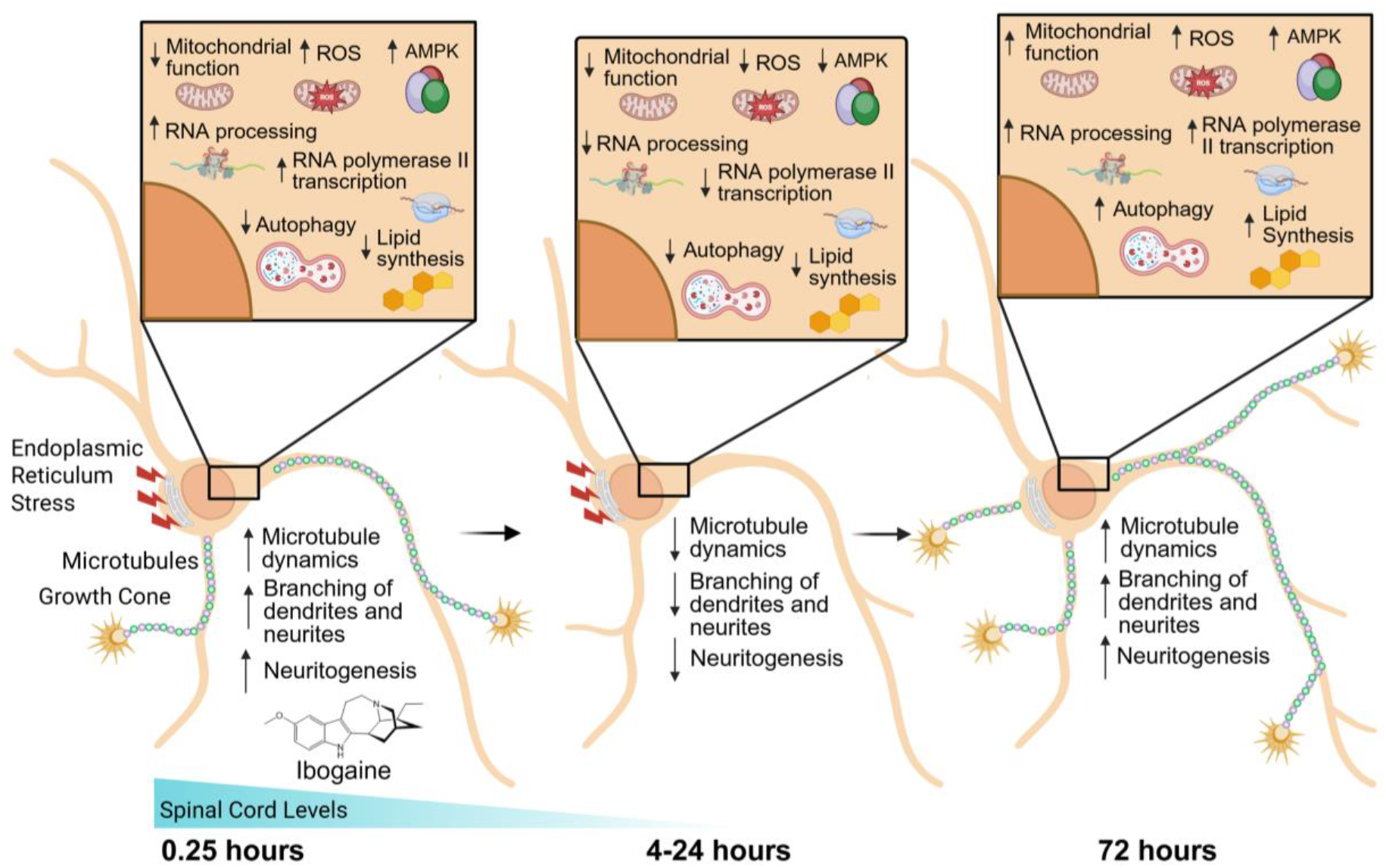

Dynamic changes in protein abundance for signaling pathways that regulate synaptic plasticity and neural repair in the spinal cord 0.25-72 hours after an oral dose of ibogaine (100 mg/kg) to female mice. Ingenuity pathway analysis revealed changes in protein abundance indicative of increased endoplasmic reticulum stress and reduced mitochondrial function, autophagy and lipid synthesis from 0.25-24 hr that return to or exceeded basal levels by 72 hr. This was accompanied by the activation, inhibition and reactivation of reactive oxygen species (ROS) production, adenosine monophosphate kinase (AMPK), microtubule dynamics, RNA processing and RNA polymerase II transcription, dendritic growth and branching and neuritogenesis signaling pathways at 0.25, 4-24 and 72 hr, respectively. These cyclic changes suggest that ibogaine re-opens a neurodevelopmental window characterized by enhanced synaptic plasticity and neural repair.

## Notes

https://data.mendeley.com/preview/c65j933r63?a=d862935e-f834-4be0-acdf-3ed6fd25a405

### Competing Interest Statement

The authors have declared no competing interest.

