## Supplementary figures and images for "Extracellular flux analysis and proteomic profiling indicate ibogaine promotes synaptic plasticity and neural repair by a metabolic preconditioning mechanism"

### Supp Fig 1.tif

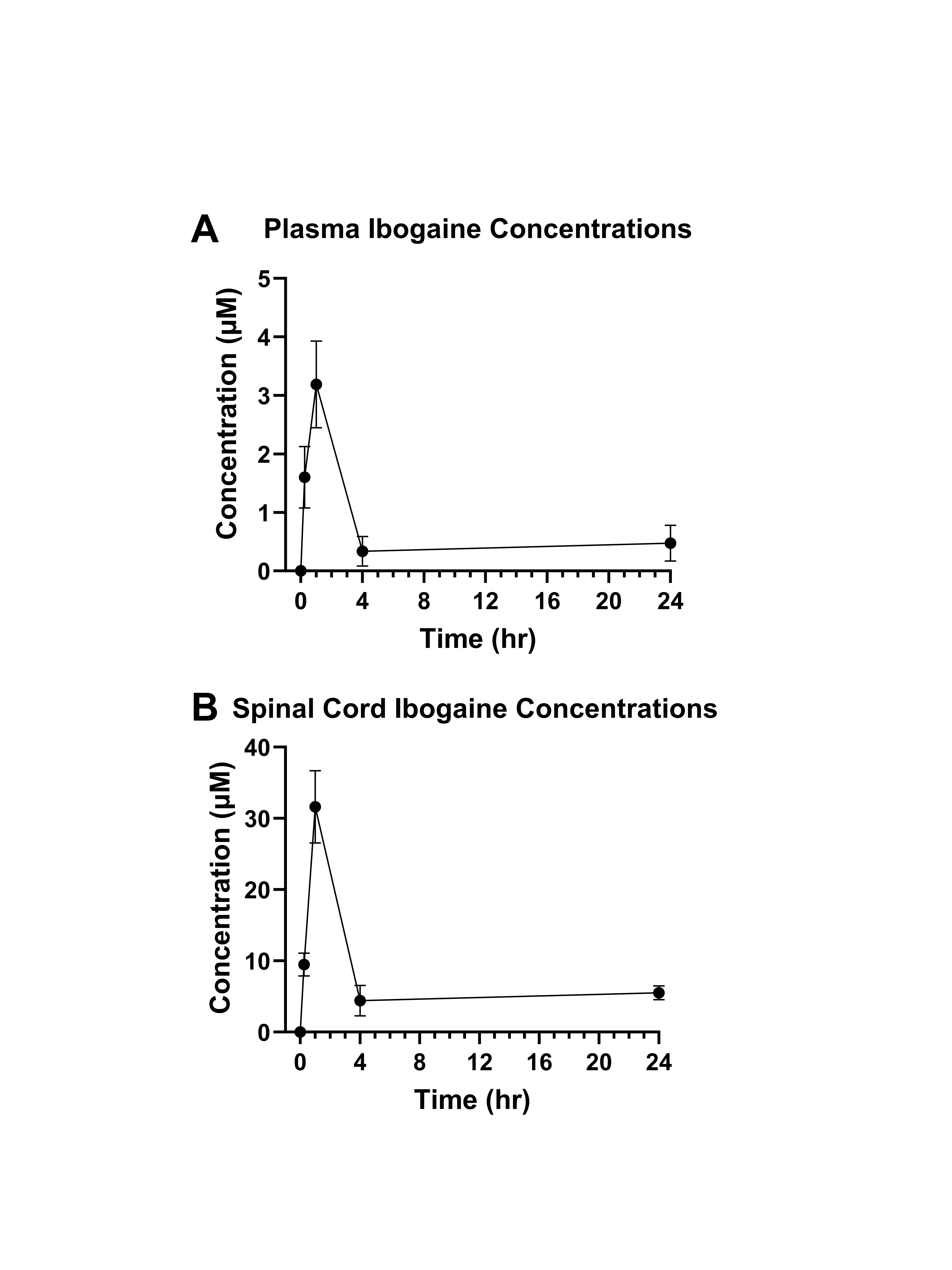

### Supp Fig 2.tif

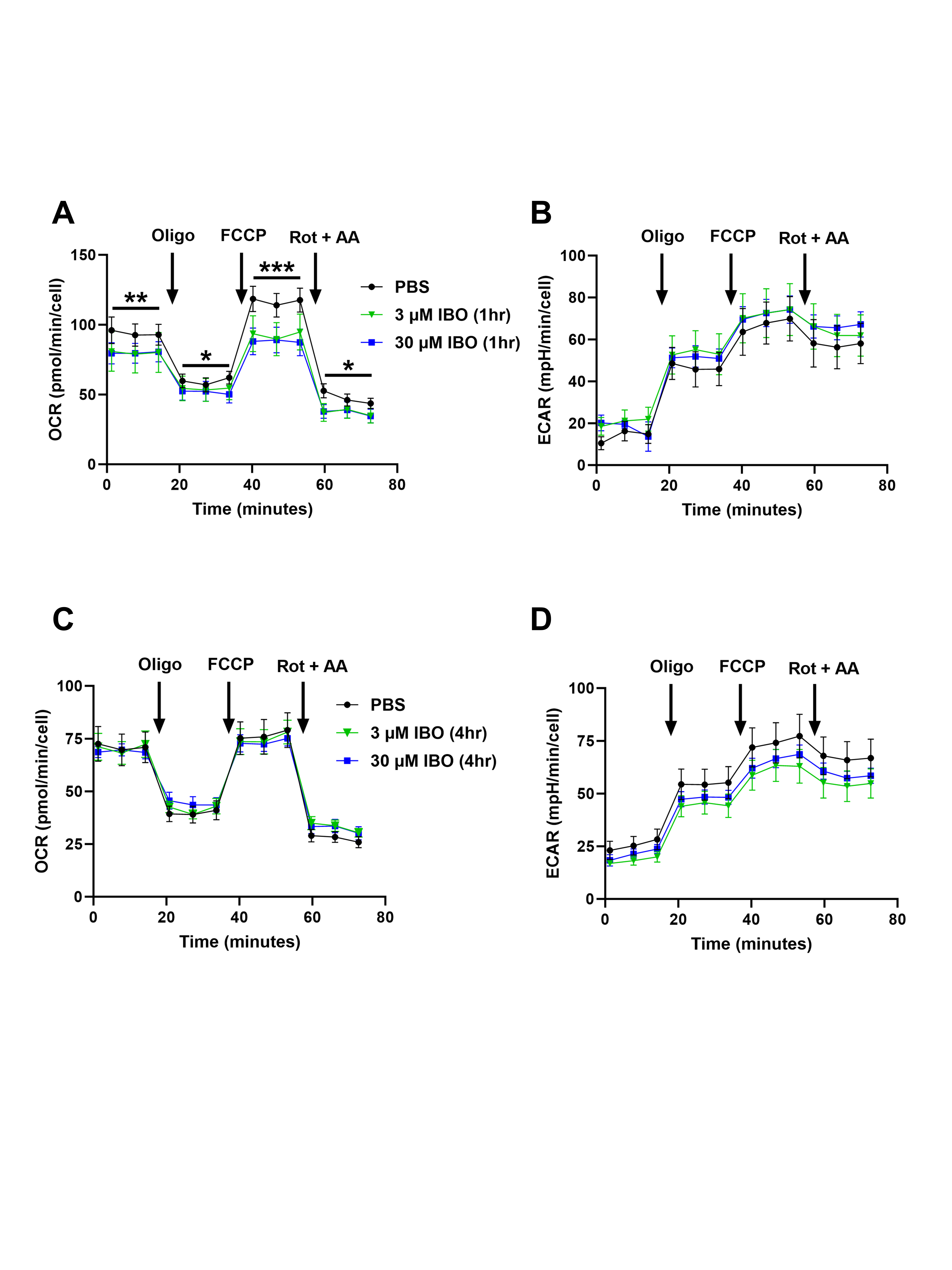

### Supp Fig 3.tif

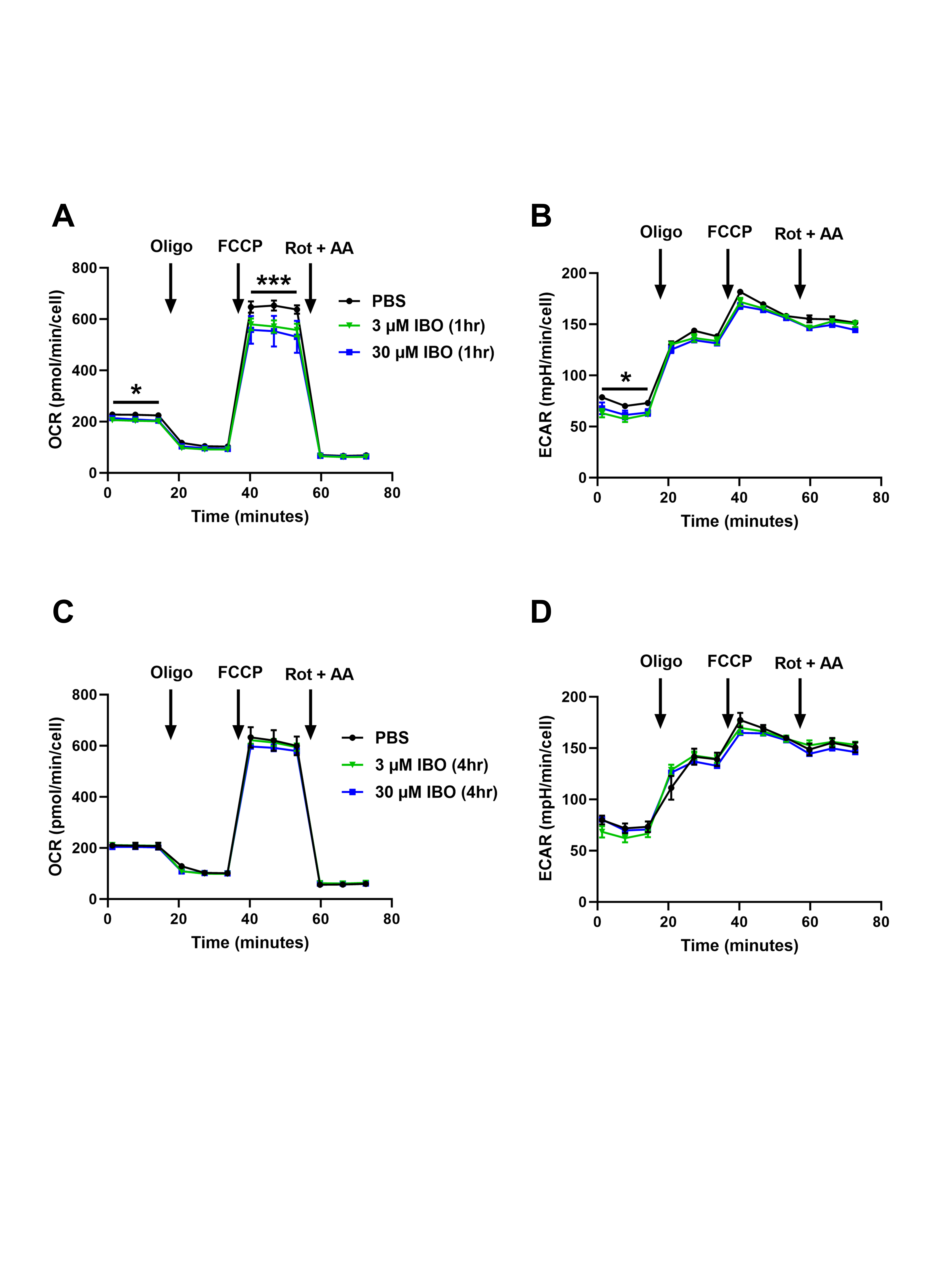

### Supp Fig 4.tif

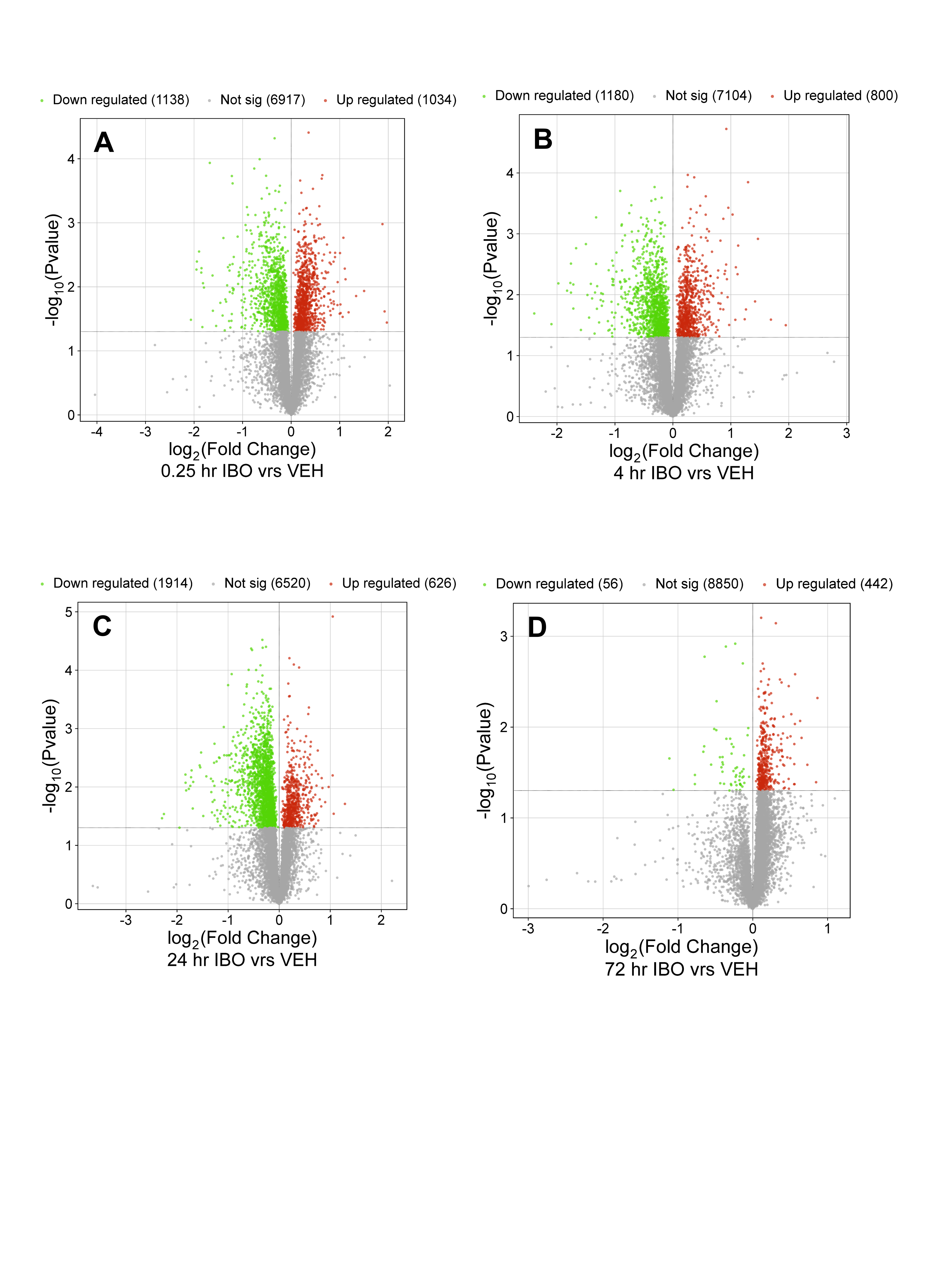
